# Rockfish: A Transformer-based Model for Accurate 5-Methylcytosine Prediction from Nanopore Sequencing

**DOI:** 10.1101/2022.11.11.513492

**Authors:** Dominik Stanojević, Zhe Li, Roger Foo, Mile Šikić

## Abstract

DNA methylation plays a crucial role in various biological processes, including cell differentiation, ageing, and cancer development. The most important methylation in mammals is 5-methylcytosine (5mC) which is present in the context of CpG dinucleotides. Sequencing methods such as whole-genome bisulfite sequencing (WGBS) successfully detect 5mC DNA modifications. However, they suffer from the serious drawbacks of short read lengths and might introduce an amplification bias. Here we present Rockfish, a deep learning algorithm that significantly improves read-level 5mC detection by using Nanopore sequencing. Compared to other methods based on Nanopore sequencing, there is an increase in the single-base accuracy and the F1 measure of up to 5% and 12%, respectively. Furthermore, Rockfish shows a high correlation with WGBS and requires lower read depth while being computationally efficient. We deem that Rockfish is broadly applicable to study 5mC methylation in diverse organisms and disease systems to yield biological insights.

## 1 Introduction

5-Methylcytosine (5mC) is among the most abundant and biologically relevant modifications involved in epigenetic regulation. On average, the frequency of 5mC in mammalian cells is 2-5% of all cytosine sites. [1]. In the human genome, 5mC is mostly present at CpG dinucleotides (where p stands for phosphodiester bond) outside CpG islands which are regions of at least 200 bp, with at least 50% GC content and 60% observed-to-expected CpG ratio [2]. Extensive in vitro and in vivo studies have shown that genomic regions with both high CpG density and high GC content tend to be hypomethylated [3]. The methylation modification is mostly symmetrical on both strands [4]. For CpG islands present at most gene promoter regions, the methylation state is typically bimodal, i.e. either fully methylated or fully unmethylated [5]. Heavy methylation in a gene’s promoter and first exon is associated with gene silencing [3], whereas intragenic 5mC is not associated with the magnitude of gene expression [5]. The methylation status of CpG island shores (which are 2 kbp regions flanking the CpG island) is highly conserved, tissue-specific and inversely related to gene expression in humans and mice [6]. On the other hand, 5mC in the gene body is more enriched in the exon and exon-intron boundary, rather than in the intron, therefore it is hypothesized that 5mC may potentially influence the alternative splicing of pre-messenger RNA [7].

In mammalian cells, DNA methylation contributes to maintaining genomic stability [8] and cellular functions [5], such as X-chromosome inactivation [9], transposon silencing [10], and genomic imprinting [11]. The landscape of DNA methylation is dynamically reprogrammed during development [12], and aberrant methylation patterns have been linked with diseases [13]. DNA methylation can also be influenced by various factors, including demographic (age, gender, race, etc.), environmental exposures (such as persistent organic/air/ heavy metal pollutants), and other risk factors (e.g. lifestyle and dietary exposures) [14]. Dynamic changes in methylation in repetitive regions are closely linked with various biological processes, such as maintaining chromosome structure and genome integrity, regulating gene transcription, embryonic development and cellular differentiation [15]. Aberrant hypomethylation in repetitive regions is biomarkers for pathological conditions from neurological diseases to cancers [16]. Therefore, it is important to profile the DNA methylation landscape to understand its cellular functions further and potentially devise therapeutic tools.

5mC was traditionally detected by affinity capture [17] and restriction enzyme digestion [18], but these techniques cannot provide the precise position of DNA modification and risk missing DNA fragments that contain only low abundance modification [19].

The gold standard method for 5mC detection at single-base resolution is bisulfite treatment [20]. Bisulfite treatment quickly converts cytosine to uracil, whereas 5mC is not influenced, and leads to differential readouts in the sequencing [21].

Upon bisulfite conversion, the sequence context readout is routinely done by either array-based methods like Illumina BeadChip platforms or next-generation sequencing (NGS) based methods.

Array-based methods are limited to a set of pre-defined selections of CpG sites in different genetic regions that influence epigenetic regulation [7]. The most comprehensive platform, Illumina MethylationEPIC microarray, contains over 850K CpG sites, holding less than 3% of the over 30 million CpG dinucleotides in the human genome [22], therefore risks missing some potentially biologically relevant CpG sites.

A combination of the bisulfite treatment and NGS, also called whole genome bisulfite sequencing (WGBS) is a popular method. However, NGS sequencing produces reads only hundreds of base pairs (bp) long. This makes their alignment unreliable in repetitive genome regions, which constitute over 66% of the human genome [23]. Bisulfite conversion of unmethylated cytosines to uracils reduces sequence complexity, influencing sequence alignment. Moreover, the strand scission and formation of abasic sites induced by the harsh bisulfite reaction condition renders up to 99.9% DNA fragments unsequenceable [24], and therefore sample information may get lost during sequencing. The DNA strands need to be amplified after bisulfite conversion to get enough material for sequencing and may introduce amplification bias.

Long-read technologies such as those of Oxford Nanopore Sequencing (ONT) and Pacific Biosciences (PacBio) enable direct sequencing of modified nucleotides which eliminates drawbacks of labour-intensive bisulfite treatment.

Methods using PacBio sequencing rely on the differences in the interpulse duration and pulse width between canonical and modified bases [25]. However, these methods are less accurate compared to the methods using ONT raw signal [26]. Moreover, CSS reads generated by PacBio sequencing are limited in length (mean length of 15 kbp or 24 kbp) and unable to bridge long repetitive regions [27]. In contrast, the ultra-long protocol in nanopore generates reads with N50 over 100 kbp [28], significantly longer than Illumina or CCS reads.

Nanopore sequencing [29] ratchets a DNA strand through the pore and measures the disruption of electrical current caused by the molecule. The raw signals obtained from nanopore sequencing have been used to detect modified bases by the differences in ionic current between modified and unmodified bases [30, 31]. There are various approaches to detect 5mC CpG modification using raw nanopore signals. They can be grouped into 3 categories of approaches: statistical testing, hidden Markov models and deep learning.

Nanoraw [32] and NanoMod [33] are tools that use statistical testing to detect modifications. They require both unmodified and modified samples. Nanoraw uses Mann–Whitney U-test combined with Fisher’s method to group neighbouring p-values. NanoMod replaces Mann-Whitney U-test and Fisher’s method with Kolmogorov–Smirnov test and Stouffer’s method, respectively. NanoRaw has been deprecated in favour of the ONT Tombo suite, which does not require an unmodified sample to detect modifications.

Nanopolish [30] and SignalAlign [34] are based on Hidden Markov Models (HMM) for detecting 5mC modifications. Nanopolish compares the likelihoods of both unmodified and modified k-mers which contain at least one CpG. If multiple CpGs are present in a k-mer, only k-mer level prediction is performed. SignalAlign uses HMM with a hierarchical Dirichlet process to learn modification effects in the raw current signal.

DeepSignal [35], DeepMod [36], and Guppy, coupled with Remora, are deep learning-based models for modification detection. While DeepMod utilizes (long short-term memory) LSTM architecture, DeepSignal combines LSTM and CNN (convolutional neural networks) architecture. DeepSignal2 replaces CNN with another LSTM, reducing computational time. Guppy is a basecaller based on recurrent neural networks (RNN) architecture that adds a modified base to the canonical alphabet and performs sequence basecalling. Recently, ONT developed a new tool named Remora to decouple modification calling from canonical basecalling. After canonical basecalling, Remora performs a second, lightweight pass through the sequence and calls modifications. Megalodon is another ONT tool built on top of Guppy (and Remora) callers. Megalodon anchors basecalling information to the reference sequence and uses called probabilities obtained from Guppy (and Remora) alongside the basecalled and reference sequence to further increase the detection performance. In the rest of the manuscript, we use the term “Megalodon” to refer to the pipeline, which includes Guppy, Remora and Megalodon. In their extensive assessment, Liu et al [37] recommended Nanopolish and Guppy for methylation analysis in the case of limited resources and Megalodon in the case of access to high-performance computing resources.

Most described methods based on nanopore sequencing require a higher coverage to correctly predict site-level methylation frequency due to their lower read-level prediction accuracy. In addition, precise read-level detection is essential in cases when particular sites in the sample are not completely methylated or unmethylated, including differences between haploids and different cell types.

Taking into account a need for a highly accurate method for read-level prediction, we set out to develop a new deep learning method using modern architecture, Transformers, for detecting 5mC. Our method, Rockfish, uses raw nanopore signal, reference sequence and alignment information to detect 5mC modification. We trained our model on both high-quality human and mouse datasets and tested it on internally sequenced H1 embryonic stem cell (H1ESc) data and a few publicly available human cancer and blood datasets. Rockfish models were extensively evaluated and compared to Megalodon and Nanopolish, the state-of-the-art modification detection tools, in the following five aspects: read-level prediction, site-level prediction, site-level correlation with WGBS, calling coverage and required running time.

## 2 Results

### Rockfish significantly improves read-level methylation prediction

Rockfish (fig. 1) predicts read-level 5mC probability for CpG sites. Rockfish consists of the signal and sequence embedding layers, a deep learning Transformer model used to obtain contextualized signal and base representation and a modification prediction head used for classification. Attention layers in Transformer learn optimal contextualized representation by directly attending to every element in the signal and reference sequence. Moreover, the attention mechanism corrects any basecalling and alignment errors by learning optimal signal-to-sequence alignment. During training, we introduced auxiliary tasks such as base prediction and signal classification tasks to further improve performance and generalization. After training the base model (teacher), we further trained a “small model” (student) using knowledge distillation to reduce the running time and improve generalization. Both models are collectively referred to as “Rockfish models” in this paper.

**Figure 1:**
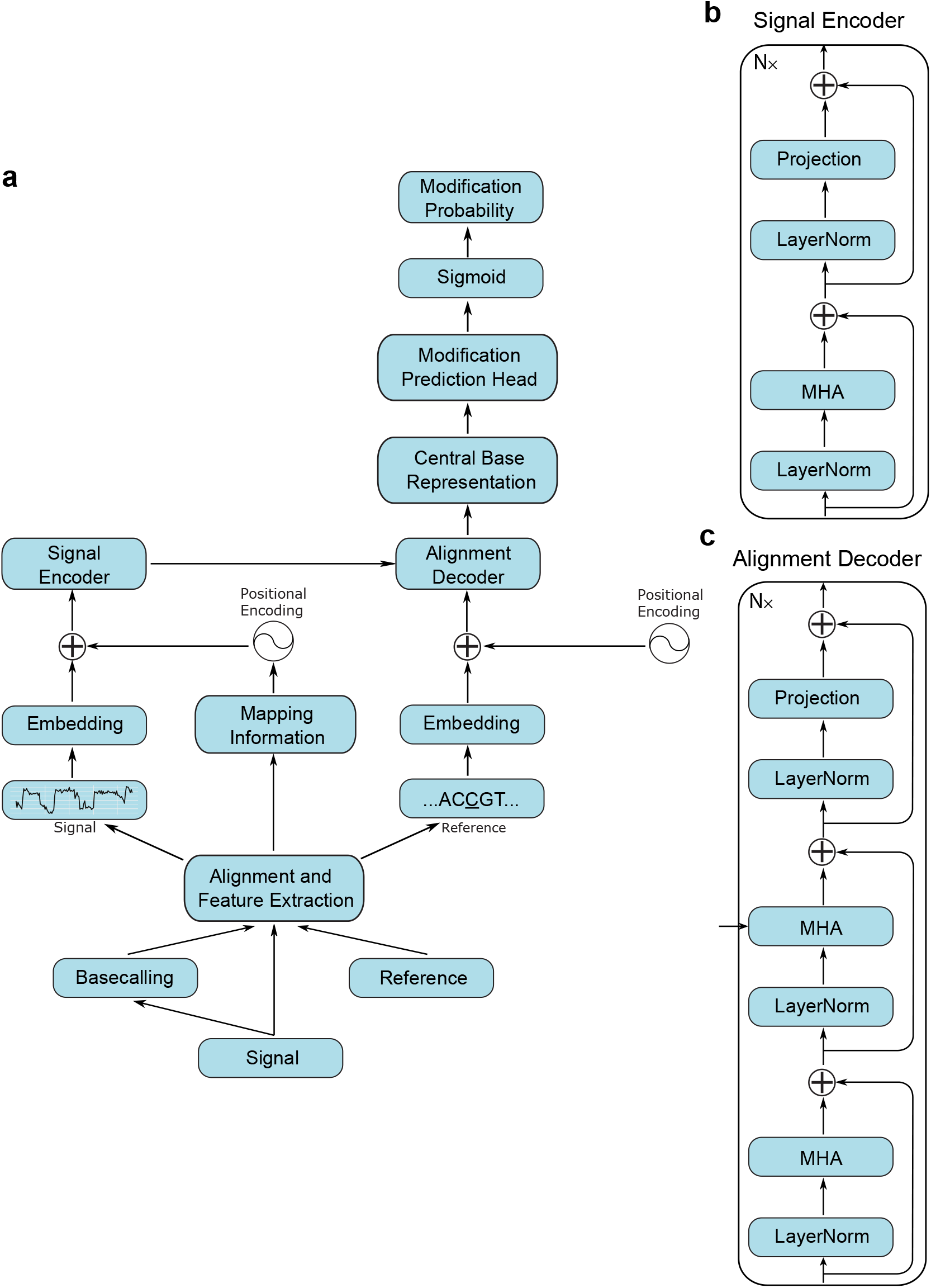
**a)** Overview of the Rockfish architecture. The nanopore signal is first processed using Guppy. Next, signal, basecalled sequence and reference are used for feature extraction. Each example consists of signal blocks, mappings and the reference subsequence. Signal blocks are embedded and processed using a signal encoder. Positional encoding for the embedded signal is given with the sine encodings and extracted mappings. Reference subsequence is embedded and fed into the alignment decoder alongside contextualized signal representation. After decoding, the representation corresponding to the central base is fed into the modification prediction head. Modification probability is obtained by applying the sigmoid function. The architecture is described in detail in the Methods section. **b), c)** Signal encoder and alignment decoder layout. They are defined as the standard Transformer (encoder-decoder) model.

Nanopore sequencing predicts methylation at a single-base, single-strand resolution. First, we evaluated Rockfish against Nanopolish and Megalodon on a read-level prediction using six different datasets. To ensure fair evaluation, we used only the read-level examples called by all ONT tools at sites covered by WGBS as the ground truth. Only fully unmethylated and fully methylated positions were used for evaluation. ONT tools are evaluated both genome-wide and in different genomic contexts (details in Methods-evaluation section): (1) singletons and non-singletons, (2) genic regions, (3) repetitive regions, (4) CpG islands, shores and shelves and (5) different GC content levels.

Read-level genome-wide results for all datasets are summarized in fig. 2(a). Both the base model and the small model significantly outperform Nanopolish and Megalodon. The base model achieves 5.15% higher mean accuracy versus the second-best ONT tool (Nanopolish for NA19240, Megalodon for rest), and the small model increases mean accuracy by 5.1%. Moreover, it can be seen that Rockfish models generalize well on the unseen chromosome (GM24385 chromosome 1), on unseen datasets corresponding to the same cell type (NA12878, NA19240 and HX1) and on unseen datasets corresponding to the different cell types (H1ESc, K562).

**Figure 2:**
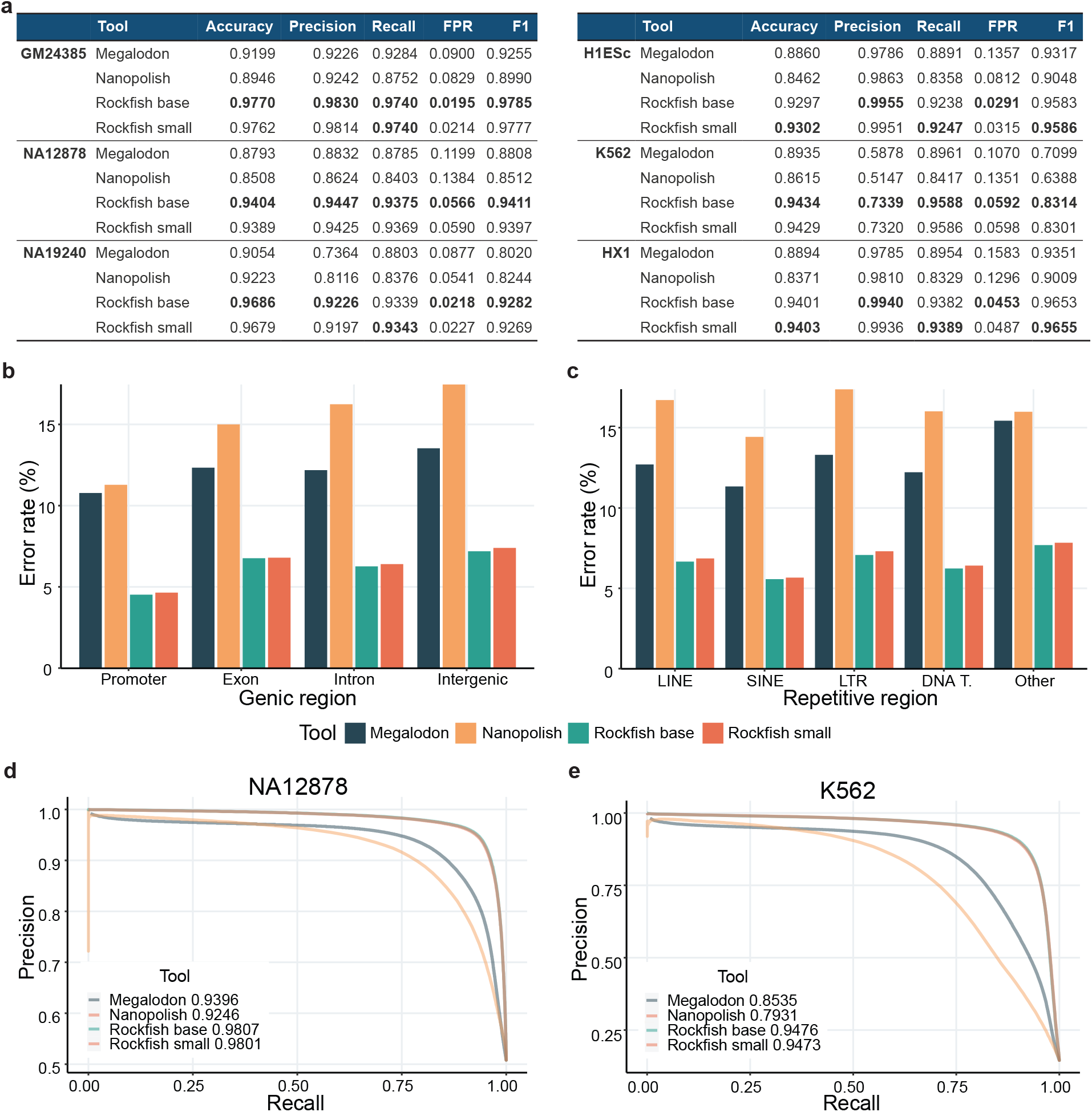
**a)** A table presenting read-level evaluation for six different datasets. For WGBS, only fully unmethylated or fully methylated sites with coverage higher than 5x were included. Rock-fish models significantly outperform Megalodon and Nanopolish on all metrics and for all six datasets. **b)** The error rate for ONT tools in different genic regions (promoters, exons, introns and intergenic regions) on the NA12878 dataset. Rockfish models significantly increase accuracy for every genic region. **c)** The error rate for ONT tools in different repetitive regions (LINE, SINE, LTR, DNA Transposons and others) on the NA12878 dataset. Rockfish models significantly increase accuracy for every repetitive region. **d)** and **e)** show precision-recall (PR) curves for NA12878 and K562 datasets that contain balanced and imbalanced read-level methylation distribution. The average precision (AP) for each tool is given in the corresponding legend. Rockfish models significantly outperform both Megalodon and Nanopolish for all probability thresholds.

Besides genome-wide evaluation, we performed read-level evaluation for various biological contexts (figs. 2(b) and 2(c), full results in Supplementary Table 1). We evaluated Rockfish for singleton (CpG sites with only one CpG up and down 10-base-pair regions),and non-singleton (CpG sites with multiple CpG sites up and down 10-base-pair regions) examples. In both scenarios, both Rockfish models outperformed Nanopolish and Megalodon. All ONT tools achieve higher accuracy for non-singleton examples since models capture information from multiple CpG sites, making prediction easier.

Figure 2(b) shows the accuracy of different ONT tools on the NA12878 dataset in different genic regions: promoters, exons, introns and intergenic regions. Both the base model and the small model outperformed other ONT tools in these genic contexts.

Figure 2(c) shows results for different types of repetitive regions: long interspersed nuclear elements (LINEs), short interspersed nuclear elements (SINEs), long terminal repeats (LTRs), DNA transposons and others. Both Rockfish models achieve higher accuracy than Nanopolish and Megalodon, and the results are consistent for all types of repetitive regions in all evaluated datasets.

Next, we evaluated Rockfish performance at CpG islands, shores and shelves. CpG shores are defined as 2 kbp regions flanking CpG islands. CpG shelves are defined as 2 kbp regions that further flank the CpG shores, upstream and downstream of CpG islands. Aberrant changes in methylation patterns in CpG islands, CpG shores and shelves are linked with cancers [38]. Rockfish models significantly outperform Nanopolish and Megalodon on all datasets (Supplementary Table 1). The Rockfish models achieve especially high accuracy at CpG islands (>97% for all datasets, >99% for GM24385).

Lastly, we evaluated Rockfish at five different GC contents: 20%, 40%, 60%, 80% and 100%. Both Rockfish models outperform Nanopolish and Megalodon for all GC contents on all datasets (Supplementary Table 1). The Rockfish models achieve higher accuracy for higher GC contents.

We then moved on to compare models’ precision and recall (PR) for different probability thresholds. Figures 2(d) and 2(e) show precision-recall curves for NA12878 and K562 datasets that contain balanced and imbalanced read-level methylation distribution. Both Rockfish models outperform Nanopolish and Megalodon for all probability thresholds. PR curves for other datasets are summarized in Supplementary Figure 1.

### Rockfish improves CpG site-level methylation prediction

After assessing Rockfish models on read-level examples, we similarly performed a site-level evaluation. First, we aggregated read-level results. For Rockfish models, we piled reads and calculated the proportion of methylated bases at a specific site without any additional filtering. Both Megalodon and Nanopolish drop uncertain predictions, resulting in a lower methylation coverage. Next, any site covered by less than 5 called reads for any ONT tool or WGBS were removed from analysis to ensure sufficient representation.

Figure fig. 3(a) shows that Rockfish small model outperforms other methods in the site level accuracy and F1 measure on all datasets apart from H1ESc, where Megalodon performs better. Results for all datasets and different genomic contexts are summarized in Supplementary Table 2.

**Figure 3:**
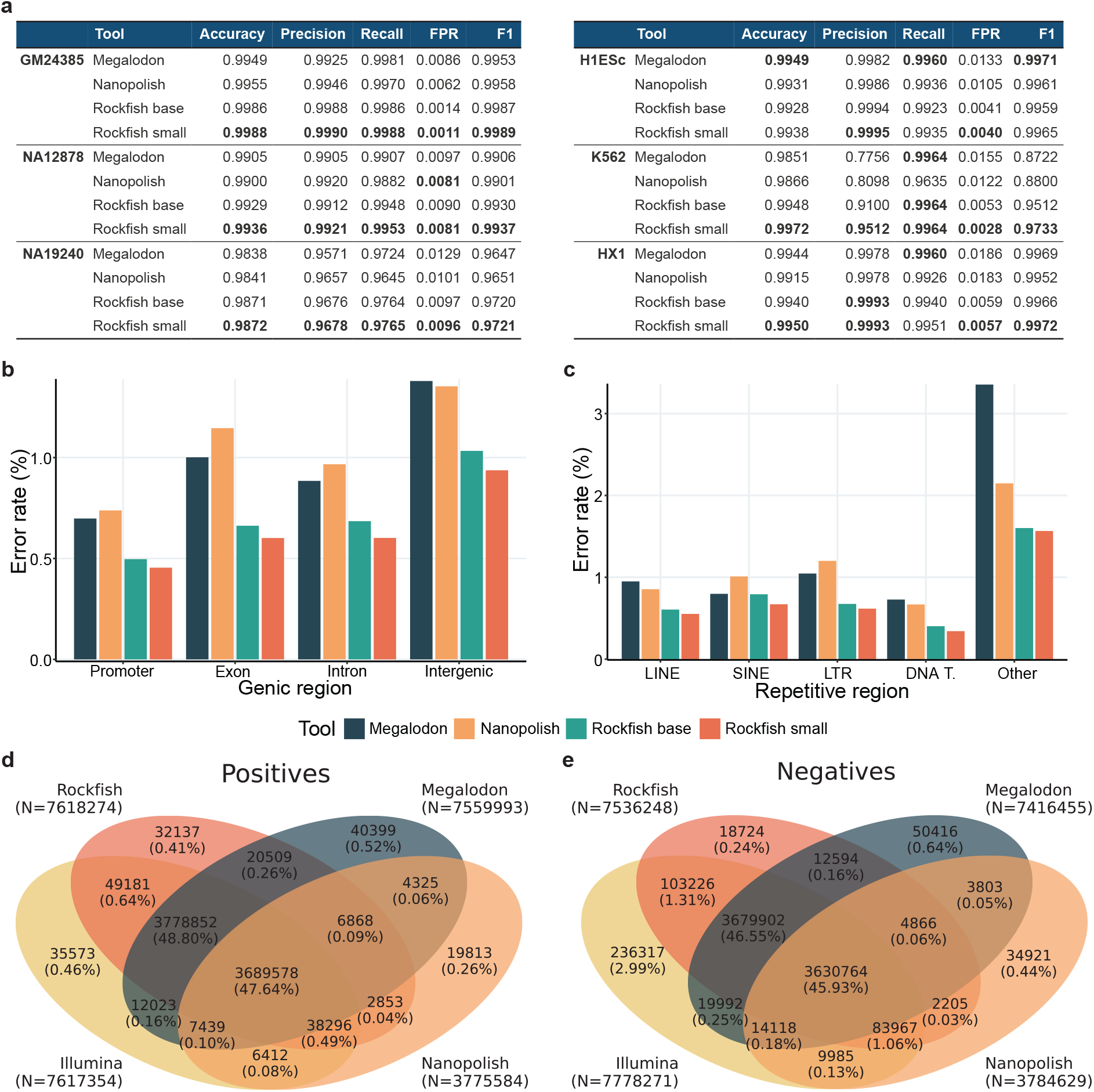
**a)** A table presenting site-level evaluation for six different datasets. Only positions with coverage higher than 5x for each ONT tool and WGBS were included. For the ground truth, we use only fully unmethylated or fully methylated positions concerning WGBS. Rockfish models outperform Megalodon and Nanopolish on most metrics and for most datasets. **b)** Error plots for different genic regions (promoters, exons, introns and intergenic regions) on the NA12878 dataset. Rockfish models reduce the error rate (1 - accuracy) for every genic region. **c)**Error plots for different repetitive regions (LINE, SINE, LTR, DNA Transposons and others) on the NA12878 dataset. Rockfish models reduce the error rate for every repetitive region. Venn diagrams representing predicted **d)** positives and **e)** negatives for different methods based on nanopore signal and the ground truth (Illumina) for the NA12878 dataset. Rockfish is represented with the small model. Sample space is defined as the set of all fully unmethylated or methylated sites called by Illumina with at least 5x. Rockfish calls the highest number of true positives and true negatives and achieves high precision and recall.

A site-level evaluation for singletons and non-singletons shows that the base model out-performs Megalodon and Nanopolish on all datasets except K562, where Nanopolish achieves slightly higher accuracy (99.70% Nanopolish vs 99.52% Rockfish base). The small model out-performs Nanopolish and Megalodon on all datasets. For singletons, base and small models achieve higher accuracy than Nanopolish and Megalodon on four datasets: GM24385, NA12878, NA19240 and K562. On H1ESc, Nanopolish (99.22%) and Megalodon (99.45%) outperform both Rockfish models (98.95% base model and 99.11% small model). On the HX1 dataset, Megalodon (99.42%) outperforms Rockfish models (99.14% base and 99.29% small).

Figure 3(b) lists error rates for different types of genic regions on the NA12878 dataset. Both Rockfish models exhibit lower error rates compared to Megalodon and Nanopolish for NA12878. Results are consistent across all datasets and genic regions, apart from HX1 (exons) and H1ESc (exons and introns), where Megalodon outperforms Rockfish models (Supplementary Table 2).

Next, we explored the performance in repetitive regions. Error rates for different types of repetitive regions for the NA12878 dataset are given in fig. 3(c). Both Rockfish models outperform Nanopolish and Megalodon, especially for type “Other”, which includes all types of repetitive regions not listed before (e.g. RNA Repeats). The same trend can be seen for all datasets, except H1ESc and HX1, where Rockfish models achieve slightly worse results compared to Megalodon and Nanopolish.

Furthermore, we evaluated Rockfish performance at CpG island, shores and shelves. Both models outperform Nanopolish and Megalodon at CpG islands and shores on all datasets. At CpG shelves, the Rockfish base model achieved the same error rate as Megalodon for the K562 ((3.49%) and H1ESc (0.54%) datasets. The small model outperforms all other models at CpG shelves.

Lastly, we evaluated the prediction performance of all ONT methods for different GC contents. Rockfish tools perform better for higher GC contents (80% and 100%) while having comparable error rates for lower GC contents with Megalodon and Nanopolish.

In the previous analysis, we evaluated only sites called by all methods. Here, we evaluate both the number of calls for and their concordance with bisulfite sequencing. Figure 3(d) describes relations between predicted positives (for methods based on ONT) and predicted positives achieved using WGBS for the NA12878 dataset. Since WGBS is currently a state-of-the-art method, in this analysis, we choose it as the ground truth. Nanopolish calls significantly fewer true positives (49.12% of all actual positives) than Rockfish (99.19%) and Megalodon (98.30%). Moreover, due to the aggressive filtering, Nanopolish calls the lowest amount of false positives (0.44% of all actual negatives). Rockfish calls a slightly lower amount of false positives (0.80%) than Megalodon (0.93%). We can conclude that Rockfish exhibits high precision while calling the highest number of sites. Figure 3(e) shows the distribution of predicted negatives with respect to bisulfite sequencing. Rockfish calls the highest number of true negatives (96.39% of all actual negatives) compared to Megalodon (94.43%) and Nanopolish (48.07%). Moreover, Rockfish calls a lower amount of false negatives (0.50% of all actual positives) compared to Megalodon (0.94%) and Nanopolish (0.60%). Similar results are achieved for GM24385 and NA19240 datasets (Supplementary Figure 2). For H1ESC and HX1 Rockfish small called more false negatives (0.70% and 0.56% of all positives respectively) compared to Megalodon (0.40% and 0.41%) and Nanopolish (0.34% and 0.38%). However, Rockfish small, also called significantly more true negatives (99.20% and 98.89% of all negatives) in comparison to Megalodon (93.38% and 90.17%) and Nanopolish (49.28% and 48.02%) for both datasets (Supplementary Figure 3). For the K562 dataset, most sites were not called (Supplementary Figure 4) due to low ONT coverage (mean coverage is 1x).

### Methylation Prediction Results Generated by Rockfish Models and WGBS are Highly Correlated

After evaluating Rockfish performance on read-level and site-level predictions, we further tested the correlation of site-level predictions with WGBS. In the subsequent experiments, we included partially methylated CpG sites to assess the concordance of methylation frequency predicted by ONT models and WGBS. Figure 4(a) demonstrates the correlation results for Megalodon, Nanopolish and both Rockfish models on all six evaluation datasets. Rockfish models outperform Megalodon and Nanopolish on all six datasets (Supplementary Table 3). Additionally, Rockfish models achieve higher correlation for most annotations, only with lower GC contents (20% and 40%) being an exception.

**Figure 4:**
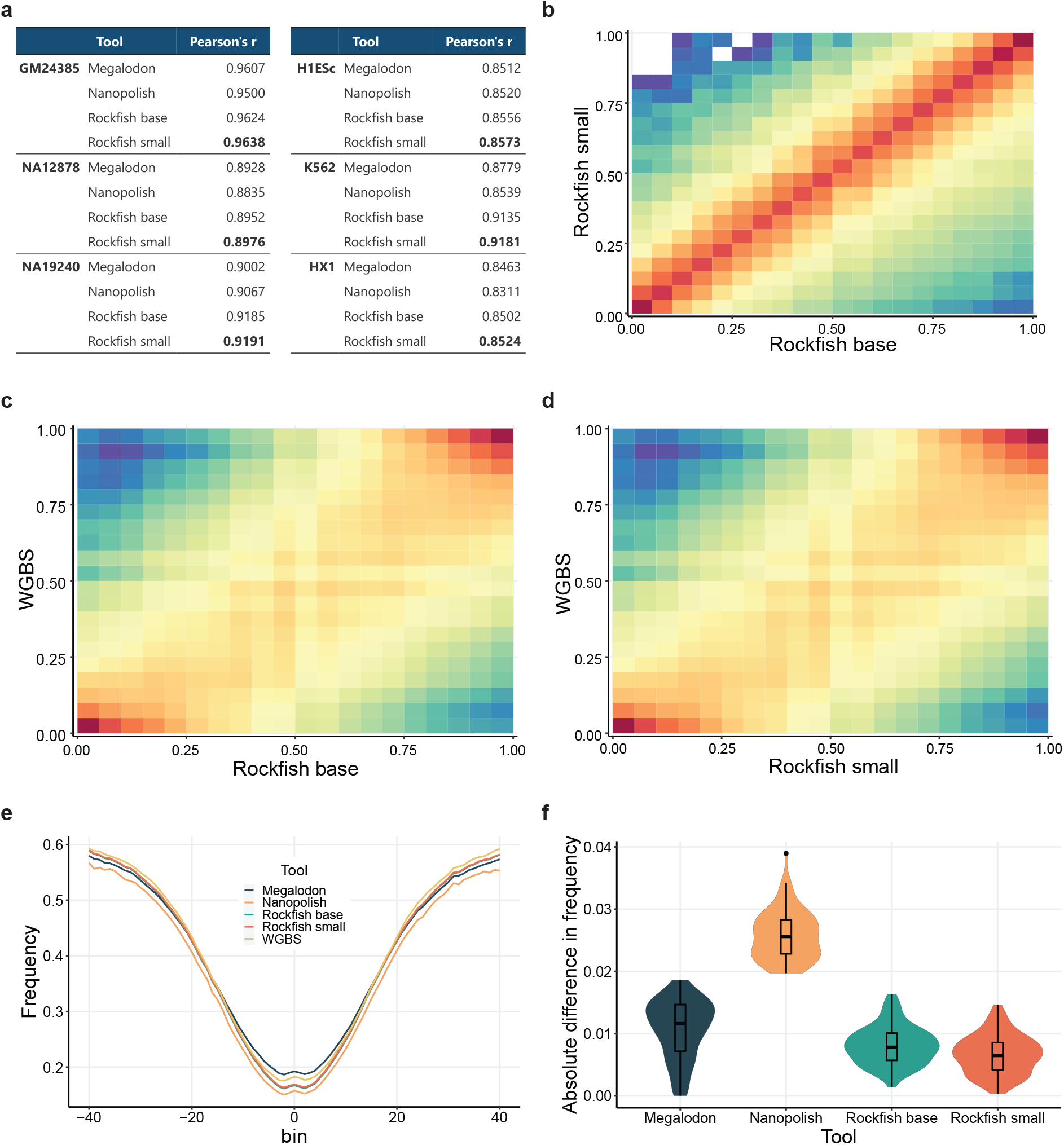
**a)** A table presenting Pearson’s correlation between the methods based on nanopore signal and WGBS. Only the sites with coverage higher than 5x for each tool were included. Both Rockfish models outperform Megalodon and Nanopolish on every evaluation dataset. **b), c), d)** 2D histograms representing correlation between **b)** Rockfish models, **c)** the base model and WGBS and **d)** the small model and WGBS for the NA12878 datasets. The models exhibit a high correlation with each other and with WGBS. Each axis is divided into 25 bins, and counts are represented on a log scale. Empty bins are represented with white colour. **e)** Methylation frequency for every ONT tool and WGBS with respect to the binned distance from the transcription start sites (TSSs) on the NA12878 dataset. Both Rockfish models show high consistency with other ONT tools and, more importantly, WGBS. **f)** the distribution of the absolute difference between every ONT tool and WGBS. Rockfish models reduce the absolute difference between ONT and WGBS.

Figures 4(b) to 4(d) show the distribution of methylation frequencies for the NA12878 dataset for (1) Rockfish base model and Rockfish small model, (2) Rockfish base model and WGBS and (3) Rockfish small model and WGBS. Methylation predictions obtained from the base and the small model exhibit a very high level of correlation (Pearson’s *r* = 0.9929, *p* = 0.0). Moreover, the results from both the base model (*r* = 0.8933, *p* = 0.0) and the small model (*r* = 0.8956, *p* = 0.0) show high degrees of correlation with that of WGBS. In addition, the results given by Rockfish models are highly correlated with that of other ONT tools, especially with Megalodon (Supplementary Figure 5). The same pattern can be seen for other evaluation datasets (Supplementary Table 3).

We then studied the distribution of methylation frequencies for the NA12878 dataset for (1) Rockfish base model and Rockfish small model (fig. 4(b)), (2) Rockfish base model and WGBS (fig. 4(c)) and (3) Rockfish small model and WGBS (fig. 4(d)). Methylation predictions obtained from the base and the small model exhibit a very high level of correlation (Pearson’s *r* = 0.9929, *p* = 0.0). Moreover, the results from both the base model (*r* = 0.8933, *p* = 0.0) and the small model (*r* = 0.8956, *p* = 0.0) show high degrees of correlation with that of WGBS. In addition, the results given by Rockfish models are highly correlated with that of other ONT tools, especially with Megalodon (Supplementary Figure 5).

Lastly, the methylation frequency was plotted with respect to the binned distance from the transcription start sites (TSSs) and the distributions of absolute differences between ONT tools (fig. 4(e)) and WGBS for the NA12878 dataset (fig. 4(f)). Results from both Rockfish models closely match that of WGBS (the median absolute difference is 0.007791 for the base model, 0.0064 for the small model), and stand lower than Megalodon (0.01168) and Nanopolish (0.02559).

### Rockfish calls more sites compared to WGBS

Next, we evaluated the number of Rockfish calls against that of WGBS for the NA12878 dataset. Complementary cumulative distribution function (CCDF) of the strand-specific calling coverage for ONT-based methods and WGBS is given in fig. 5(a). The calling coverage is defined with respect to the outputs given by each tool. Since Rockfish does not perform any filtering of the read-level examples, it is able to call more CpG sites than Megalodon and Nanopolish for coverages lower than the sequencing coverage. Besides, Rockfish achieves the highest mean strand-specific calling coverage (∼17x) compared to other ONT-based methods (Megalodon ∼14x, Nanopolish ∼16x). Although WGBS was sequenced at much higher strand-specific coverage (∼48x vs ∼23x ONT), the mean strand-specific calling coverage (∼18x) is close to that of Rockfish.

**Figure 5:**
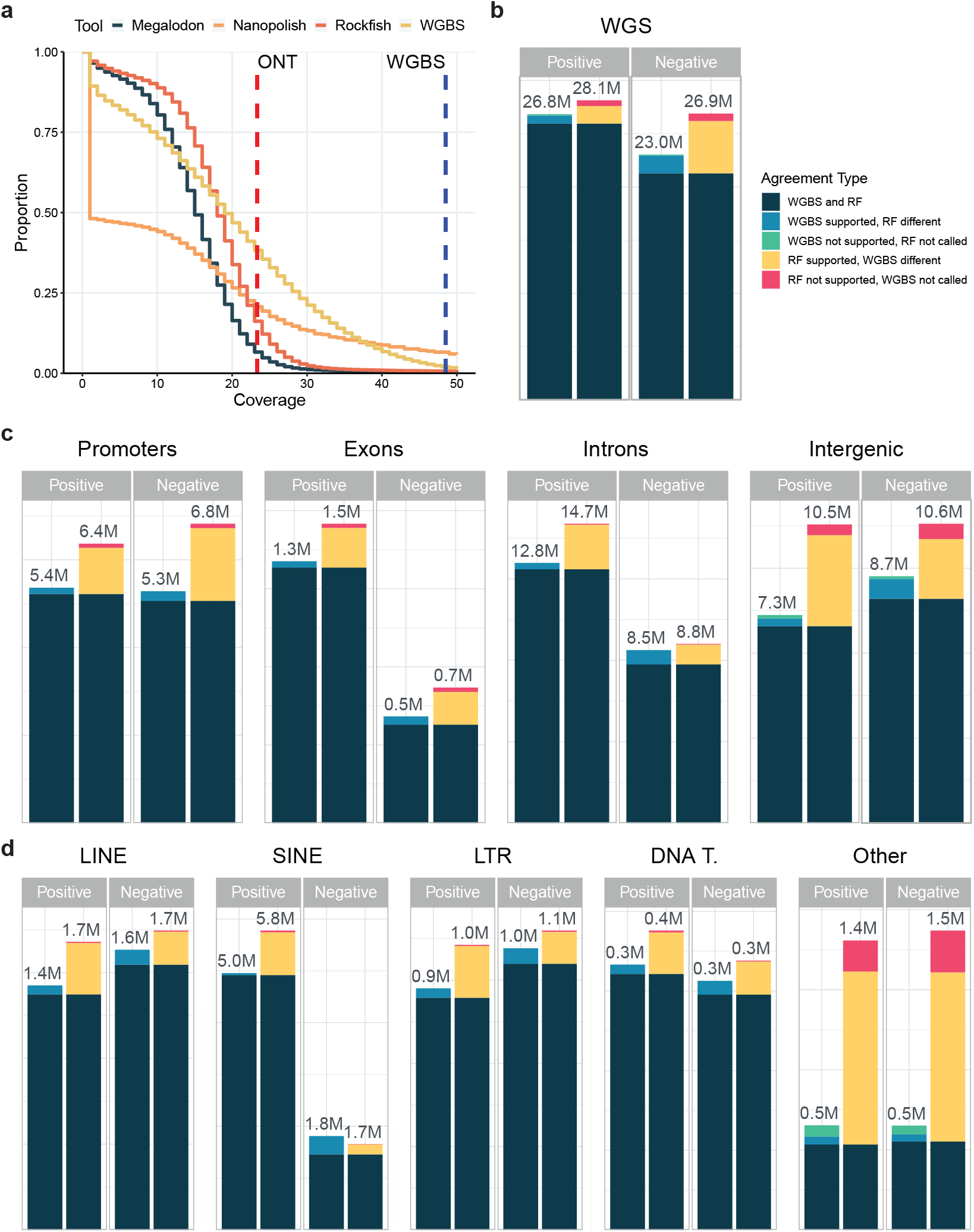
**a)** Complementary cumulative distribution function (CCDF) of the strand-specific calling coverage for each ONT-based method and WGBS for the NA12878 dataset. Vertical lines represent strand-specific sequencing coverage for ONT (∼23x; red) and WGBS (∼48x; blue). **b)** Stacked bar charts representing counts of highly confident positive and negative sites for WGBS (left) and Rockfish small (right; RF) for the NA12878 dataset. A CpG site is defined as positive if the coverage is at least 5x and methylation frequency is at least 50%. A CpG site is defined as negative if the coverage is at least 5x and the frequency is less or equal to 50%. Sites with coverage less than 5x are labelled as not called. We define three categories of highly confident sites: (1) WGBS and Rockfish are concordant, (2) WGBS and Rockfish calls differ with the target tool having support from at least one other ONT-based method, (3) WGBS without support and Rockfish without call and vice-versa. The numbers are given in millions (10^6^). Rockfish calls more highly confident sites than WGBS on the whole genome. **c)** Stacked bar charts representing counts of highly confident positive and negative sites for different genic and intergenic regions. Rockfish calls more highly confident sites for all genic and intergenic types. **d)** Stacked bar charts representing counts of highly confident positive and negative sites for different repetitive regions. Except for unmethylated sites in SINE, Rockfish calls more highly confident sites than WGBS.

Moreover, we compared the number of highly confident positive and negative calls for WGBS and Rockfish. Highly confident calls were divided into three categories. The first category corresponds to the CpG sites for which both Rockfish and WGBS made concordant calls (both methods called a CpG site to be either positive or negative). In the second category are the positions for which the calls for WGBS and Rockfish differ with the target method (either WGBS or Rockfish) having support from other methods (Megalodon and/or Nanopolish). The last category is the unique sites for which the target tool does not have any support and the other tool does not produce a call. A CpG site is called positive if it is covered by at least five reads with a methylation frequency higher than 50%. A CpG site is called negative if it is covered by at least five reads with a frequency less or equal to 50%. If a site was covered with less than five reads, it was deemed as uncalled. The results on the whole genome for NA12878 are given in fig. 5(b). Rockfish calls more highly confident positives and negatives (28.14M and 26.90M) compared to WGBS (26.84M and 23.05M). The majority of the calls for both WGBS and Rockfish were concordant calls (25.95M for positives, 21.28M for negatives). Rockfish had more support for both positive sites (1.65M) and negative sites (4.92M) compared to WGBS (0.75M and 1.66M, respectively). Lastly, Rockfish had more unique, unsupported calls (0.54M and 0.70M) compared to WGBS (0.13M and 0.10M). We also explored the number of highly confident calls in genic and intergenic regions (fig. 5(c)) and repetitive regions (fig. 5(d))). Except for SINE, where WGBS calls more negatives (1.81M vs 1.66M), Rockfish calls more highly confident positives and negatives for all types of annotations. The biggest difference between Rockfish and WGBS occurs in intergenic regions (Rockfish calls 43.55% more positives and 21.29% more negatives) and for “Other” repeat types (Rockfish calls 176.48% more positives and 187.14% more negatives). Full counts are given in Supplementary Table 4.

### Running time required by Rockfish models is comparable with that of other ONT tools

Besides the high prediction quality, another important aspect of a high-quality bioinformatics tool is the running time. In this section, we assessed the running time of all ONT tools. The K562 dataset was used to evaluate the running time of each ONT tool. The dataset contains 263019 reads stored in single read FAST5 format. Megalodon calls methylation in an end-to-end fashion (basecalling, alignment and methylation calling), so only one command is run. Nanopolish pipeline requires running four sequential commands: basecalling, indexing, alignment and methylation calling. Rockfish currently requires running two consecutive commands: basecalling and methylation calling. The mean running time for all commands used for every ONT tool is given in fig. 6 and in Supplementary Table 5. As expected, Megalodon is the fastest tool (mean running time of 4397 s) with a running time similar to Guppy canonical basecalling (4458 s). The Rockfish base model is the slowest (8469 s), being slightly slower than Nanopolish (7660 s). The Rockfish small model reduces methylation calling time by 58.42% compared to the base model for the K562 dataset. In the pipeline, which included the small model, Guppy was the bottleneck taking 72.77% of the total running time.

**Figure 6:**
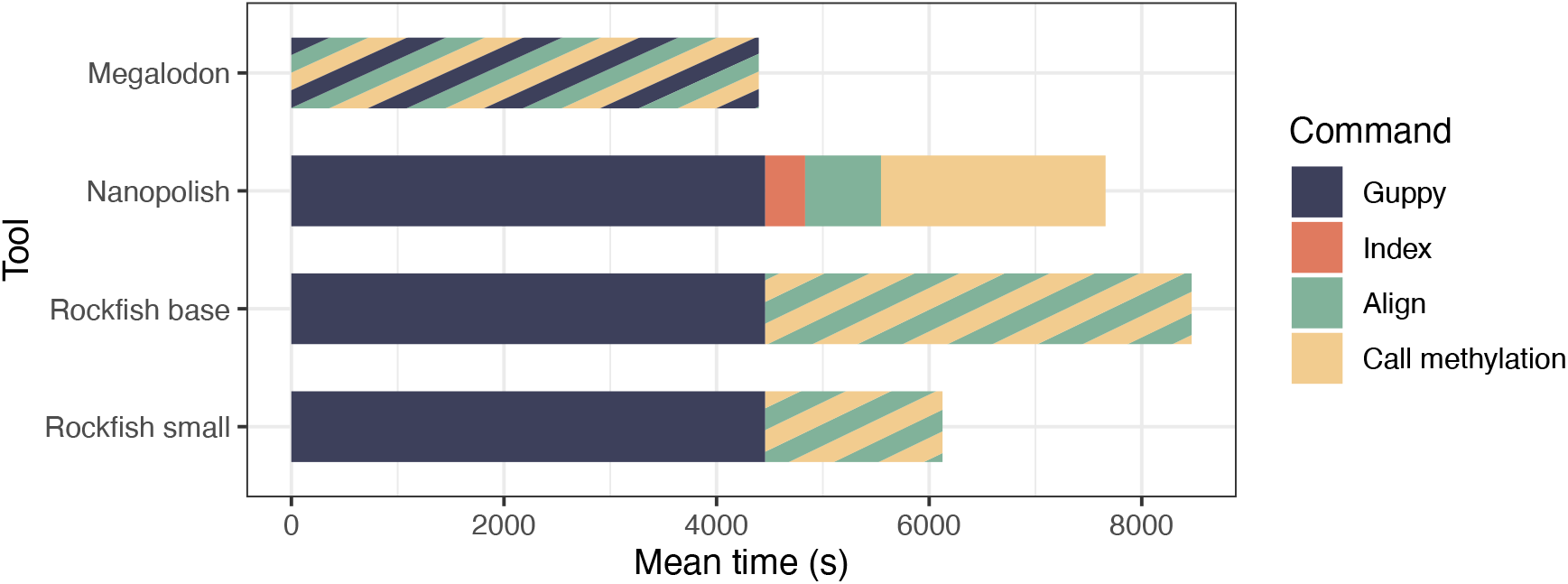
Mean running time for each ONT tool on the K562 dataset. Megalodon requires invoking only one command which includes basecalling, alignment and methylation calling. Rockfish requires invoking two commands -basecalling and inference. The inference consists of both alignment and methylation calling. Nanopolish requires invoking four commands. Different command types are plotted in different colours. Each command is called three times.

## 3 Discussion

In this paper, we present Rockfish, a deep-learning method for detecting 5mC at CpG sites in DNA at single-base, single-strand resolution. It utilizes a Transformer network, an architecture used in various visual, language and speech recognition tasks. Rockfish models do not rely on the explicit re-segmentation step. Rather, the signal information is combined with the reference subsequence and alignment information to obtain local alignment using multi-head attention. In addition, we introduced a few auxiliary tasks, including base masking and signal classification, to further increase the generalization performance. Finally, to reduce the computation time without sacrificing the generalization performance, we used knowledge distillation on the partially methylated sites in the human genome and unlabelled mouse data.

We systematically compared the performance of Rockfish models against Megalodon and Nanopolish, state-of-the-art methods for detecting 5mC, from four aspects: read-level prediction, site-level prediction, site-level correlation with WGBS, and running time required.

For read-level evaluation, Rockfish models significantly outperformed Megalodon and Nanopolish on all six internally-generated and open-access datasets from different biological contexts. There is an increase in accuracy and F1 measure of up to 5% and 12%, respectively. Precision-recall curves show that the Rockfish models are resistant to class imbalance. Furthermore, the models outperform Megalodon and Nanopolish for all probability thresholds. Significantly higher precision on the read-level prediction lowers the required coverage depth and reduces the costs for profiling the methylation landscape. Accurate read-level methylation prediction in long reads is crucial for the haplotype phasing [39], and aids haplotype-aware diploid assemblies [40]. Phasing facilitates potential applications such as cell type deconvolution in heterogeneous samples (e.g. blood or tumour samples [41]), as well as detecting allele-specific methylation typical for imprinting [42] and chromosome X inactivation [43] in homogenous samples.

Next, for site-level predictions, Rockfish models reduced the error rate for most datasets and most biological contexts while including all called examples from the read-level prediction. Rockfish models performed slightly worse only on predicting methylation status in singletons and in regions with lower GC contents (20%, 40%). Inferior performance for singleton context may be due to the training dataset GM24385. RRBS was used to generate the ground truth for the GM24385 dataset. RRBS first digests the genome with restriction enzymes prior to bisulfite treatment. Due to restriction enzyme preference and size selection, the library distribution may be skewed towards non-singleton and CpG-rich areas. Even though Rockfish performed slightly worse in regions with lower GC content compared to other ONT tools, CpG instances from these regions are the minority in the human genome. Of all human CpG instances, only 1.29% are from the 20% GC content region, while 17.79% are from the 40% GC content region in the CHM13 reference genome, therefore the impact is not detrimental for overall detection. Adding more singleton training examples may help further improve the performance of Rockfish models.

Both Rockfish models demonstrated a higher correlation with WGBS compared to Megalodon and Nanopolish. ONT-based methods had a smaller difference between sequencing and calling coverage than WGBS, possibly due to alignment ambiguity typical for short-read sequencing. Moreover, Rockfish had the highest mean strand-specific coverage compared to the other ONT tools and exhibited a higher number of highly confident calls than WGBS, especially for intergenic sites and “Other” - repetitive regions excluding SINE, LINE, LTR and DNA transposons. Lastly, the running time that Rockfish models required were comparable to that of Megalodon and Nanopolish.

The presented results demonstrate that Rockfish is a powerful and reliable method for extracting methylation information from the ONT raw signals. Further, the small model outperformed the base model on all datasets and required a shorter running time, showing the benefit of additional data and knowledge distillation.

5mC modifications are enriched in various genic elements, and further relate to many biological phenomena, such as transcription regulation [44], chromatin architecture [45], diseases [46], ageing [47], memory formation [48], exercise [49] and many more. Therefore, the ability to detect 5mC modification at a single-base, single-strand resolution is critical for a deep understanding of the role DNA methylation plays in these biological phenomena. This knowledge might contribute to the early detection of disease onset, as well as patient stratification, treatment strategy choice, and in the future even epigenome editing as a new direction of therapeutic targets.

Finally, due to its architecture, the Rockfish pipeline might be easily adapted to detect various types of DNA and RNA modifications.

## 4 Methods

### 4.1 Feature Extraction

Sequenced FAST5 files are first processed using Guppy basecaller^1^. The Guppy version used for all experiments is 5.0.14. We used a super-accurate DNA basecalling model since it achieves the highest basecalling accuracy. We use an additional *--fast5 out* flag to output the computed move table into a FAST5 file. The move table defines sequence-to-signal alignment that is used for obtaining final reference-to-signal alignment, similarly to Megalodon.

After obtaining sequence-to-signal alignment, basecalled ONT reads are aligned to a reference genome using mappy, minimap2 biding for Python [50]. The Mappy version used for the sequence alignment is 2.24. The reference genome used for human datasets is T2T-CHM13 (v2.0), the complete assembly of a human genome [51]. We use Genome Reference Consortium Mouse Build 39 (GRCm39) for mouse datasets. We choose the best primary alignment and generate reference-to-query mapping by parsing the CIGAR string. Unmapped reads are not processed further.

Next, we find relevant positions - i.e. CpG dinucleotides in the reference genome and extract a subsequence of length l = 31 around each relevant position. For each subsequence, we extract corresponding p signal points. We divide signal into signal blocks **B** ∈ ℝ^*s*×*b*^ where each block corresponds to b points, *s* = *p*/*b*. The parameter b is defined by a specific Guppy model. For quality and computational reasons, examples with less than s or more than 4s blocks are discarded. Moreover, we extract relative reference indices 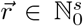 and relative query indices 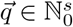. Both 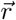 and 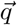 will be later used as positional encodings.

### 4.2 Architecture

The Rockfish architecture consists of four components: signal embedding layer, reference embedding layer, Transformer and modification prediction head. First, we use a projection layer **W**^*S*^ ∈ ℝ^*b*×*f*^ to embed signal blocks into a local representation **S** ∈ ℝ^*s*×*f*^, **S** = **BW**^*S*^. Here parameter f is the dimension of both signal and reference latent space.

Next, we add positional encodings to the signal representation since the Transformer architecture itself is not processing data sequentially. We add four types of positional encoding: both cosine and sine encodings of the absolute position, the cosine of the relative reference indices and the cosine of the relative query indices. Alongside traditionally used cosine and sine encodings [52], we decided to use both reference and query indices to provide additional positional information to the model. The final positional encoding matrix **P** ∈ ℝ^*s*×*f*^ for a specific example is given with:

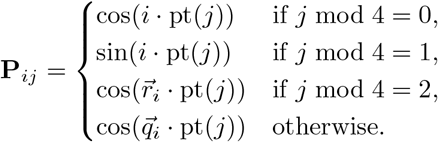

Positional term function *pt* is defined as pt(*i*) = 10000^−4⌊*i/*4⌋*/d*^. The final input to the Transformer encoder is obtained by adding positional encodings *P* to the signal representation *S*.

We use a Transformer encoder [52] to capture contextual information between the signal points by iteratively updating signal representation **S**. To improve training stability, the original (Post-LN) encoder layer is replaced with the Pre-LN layer [53]. First, inputs to the encoder layer are normalized using layer normalization [54]. Next, we update the signal representation by using multi-head attention (MHA):

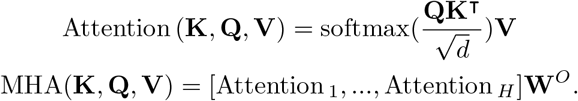

where [$#x00B7;] is the concatenation operator, H is the number of heads, d = f/H and **W**^*O*^ ∈ ℝ^*f* ×*f*^ is an output projection matrix. Scaled dot-product attention for each head is given as Attention_*i*_ = Attention 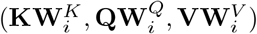 where 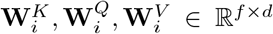 are projection matrices. Finally, the result of the multi-head attention is added to the input.

In encoder multi-head attention is defined as self-attention since matrices **K, Q** and **V** all correspond to the normalized input. After multi-head attention, signal representation is again normalized using layer normalization and fed to the linear projection layer. Linear projection layer is a simple two-layer feed-forward network given as: Projection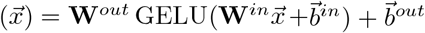 where GELU is a Gaussian Error Linear Unit [55], W^*in*^ ∈ ℝ^*f* ×*p*^, 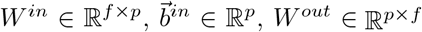, W^*out*^ ∈ ℝ^*p*×*f*^ and 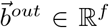. The final output is given by adding projection output to the projection input. This process is repeated *L*_*enc*_ times.

After obtaining contextualized signal representation, we embed reference bases using simple look-up table **E**^*B*^ ∈ ℝ^6×*f*^ to obtain localized sequence representation **R** ∈ ℝ^*l*×*f*^. Except for four canonical bases, we also define two extra tokens: unknown token and mask token. Unknown token [UNK] represents all non-canonical bases with respect to the FASTA format. Mask token [MASK] is used for during training for base prediction task. Next, positional encodings (same as in [52]) are added to the localized sequence representation.

We use the Pre-LN Transformer decoder to obtain contextualized sequence representation. In the decoder, the starting sequence representation is iteratively updated using decoder layers. Each layer uses both current sequence representation and contextualized signal representation to update the sequence representation. First, the current representation is normalized by applying layer normalization. Next, the representation update is performed using self-attention, the same as in the encoder layer. The output from self-attention is added to the input. The resulting representation is then normalized and passed to the multi-head attention where query matrix **Q** corresponds to the current representation and matrices **K** and **V** correspond to the contextualized signal representation. The motivation for using multi-head attention is to learn the alignment between signal and reference sequence and to update sequence representation with relevant signal information. Next, MHA output is added to the input, normalized and then passed through the projection layer and added to the input, the same as in the encoder. We repeat this process L_*dec*_ times to obtain the final contextualized sequence representation. More details regarding Transformer architecture and the pseudocode can be found in [56].

To obtain modification probability, we take the contextualized representation corresponding to the central cytosine 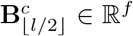 and pass it through the modification prediction head. The modification prediction head is a linear layer which outputs unnormalized modification probability.

### 4.3 Training and evaluation

#### 4.3.1 Modification prediction task

We model modification prediction as a binary classification task. Loss for i-th example is given with: 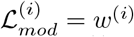. BCE(*z*^(*i*)^, *y*^(*i*)^) where BCE is binary cross-entropy loss, z^(*i*)^ is unnormalized probability, y^(*i*)^ is the ground truth and w^(*i*)^ is the singleton weight for i-th example. We add hyperparameter w to the loss to force the model to focus on singleton examples since the prediction is harder for them. Singleton examples are defined as examples with only one CpG dinucleotide in the reference subsequence. For singletons w > 1 and for non-singletons w = 1.

#### 4.3.2 Auxiliary tasks

To improve learning and generalization, we implement two auxiliary tasks during training: base prediction task and signal classification task. Both base prediction and signal classification tasks are related to masking, a self-supervision technique used during model pre-training [57][58] or model training [59]. In every iteration, for the base prediction task, we randomly choose *p*_*mask*_ of all reference bases and mask them using the mask token. Moreover, we also randomly flip *p*_*flip*_ of all bases and choose the new base with equal probability. During training, contextualized representations corresponding to the masked and flipped positions are passed to the base prediction head to obtain logits for each base. These logits will be used to predict the correct bases with cross-entropy loss ℒ_*bases*_. The base prediction task forces the model to learn to predict the correct reference bases. This helps the model to learn the local signal-to-reference alignment and correct any errors introduced during sequencing, basecalling and alignment. Moreover, it helps the model to learn the alignment between the signal and reference sequence.

The second auxiliary task is the signal classification task. The idea of the signal classification task is to force the model to learn the context for each signal block independent of the specific task. Since signal blocks are continuous, we introduce representative vectors, named codewords, to be used for classification. The collection of codewords is a codebook *C* ∈ ℝ^*f* ×*K*^, K being the number of different codewords. First, we randomly choose *p*_*signal*_ of all the signal blocks *S* that will be masked. Next, we calculate the probability of a local signal representation belonging to i-th class: 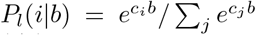.Target class for a specific signal block is given with *t* = *argmax*_*i*_*P*_*l*_(*i*|*b*). Elements of masked blocks, relative reference and query indices are all set to zero. After a forward pass through the Transformer encoder, we calculate the probability of contextualized signal representation belonging to *i*-th class in the same way as above and use these probabilities as predictions for cross-entropy loss ℒ^*signal*^. To ensure non-trivial solution, we introduce diversity loss 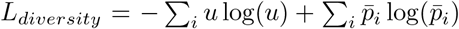 where u = 1/K and 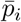 is the average hard probability of i-th class calculated for each batch.

#### 4.3.3 Final loss and optimization

The total loss for the training is given as a linear combination of all losses:

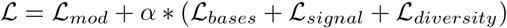

where α is a scaling parameter. All weights are optimized using a modified version of Adam [60] which decouples weight decay from the optimization procedure [61]. In our experiments, the learning rate is set to 3*e*^−4^ and weight decay is set to 1*e*^−4^. Running average coefficients were set at their default values *β*_1_ = 0.9, *β*_*2*_ = 0.999.

#### 4.3.4 Evaluation

We performed read-level, site-level and correlation evaluations to compare our method against other ONT methods. For read-level and site-level we reported accuracy, precision, recall, falsepositive rate (FPR) and F1 score for each ONT tool. Moreover, we plotted the precision-recall curve for read-level predictions. Venn diagrams were used to describe the relations between site-level predictions for all ONT methods and WGBS. All evaluation metrics were calculated using their standard definitions. Average precision was calculated as defined in [62]. The evaluation was performed on chromosomes 1-22, X and Y if the data corresponds to the male genome.

We used WGBS (or RRBS) data as the ground truth. The pipeline used for processing WGBS data includes Trim Galore ^2^ used for adapter and quality trimming and Bismark [63] used for alignment, deduplication and methylation extraction. The pipeline is fully described in the Supplementary Material. To reduce bias related to the misaligned Illumina reads we defined lower and upper coverage bounds. For read-level and site-level evaluation, we excluded genomic positions which have coverage less than max(*P*_5_, 5) where *P*_5_ is the 5-th percentile for coverage distribution in the corresponding WGBS. For correlation evaluation, we also put an upper bound to be *P*_95_ (95-th percentile). Moreover, for read-level and site-level evaluation, we removed all positions that are partially methylated. A CpG site is defined to be partially methylated if the frequency in bisulfite sequencing is between 0.01 and 0.99.

For read-level evaluation, we evaluated only examples that are present in all ONT tools. For site-level and correlation evaluation, we used only positions with ONT coverage higher than 5x. We reported read-level, site-level and correlation results for different types of annotations: gene annotations, repetitive regions, CpG islands, GC content, and CpG count, similarly as in [37].

For gene annotations, we defined four categories: promoters, exons, introns and intergenic positions. Annotations for transcription start sites (TSS), genes and exons were extracted from the annotation file. Examples were labelled as promoters if they belong to a region ±2000 around the TSS. Positions corresponding to the introns were obtained by subtracting exons from genes using *intersectBed* [64]. All other positions were labelled as intergenic positions. If a position was labelled with multiple annotations, we defined the following precedents: promoter > exon > intron > intergenic.

Annotations for repetitive regions were generated using RepeatMasker. We defined five repeat categories: SINE, LINE, LTR, DNA transposons and “Other” (positions belonging to Simple, Low complexity, Satellite, RNA, Others or Unknown repeat class).

For the analysis of CpG islands, we defined three categories: CpG islands, CpG shores and CpG shelves. GC content corresponds to the frequency of cytosines and guanines in 5-base windows. For CpG count we defined two categories: singleton and non-singleton CpG. The positions were labelled as singletons if there was only one CpG (central) in the 25-bp region around the central CpG. Otherwise, a position was labelled as a non-singleton.

Furthermore, we plotted methylation frequencies with respect to the binned distance to TSS for the NA12878 dataset. For each TSS, we calculated the corresponding bin according to bin = (*pos* − *P* + ⌊*B*/2⌋)/B where pos is the tested position, *P* is the TSS and *B* is the bin size. For TSS evaluation we set bin size to *B* = 50 base-pairs, same as in [37]. In these experiments, we did not perform intersection between tools, but we filtered out positions individually. Positions were discarded if coverage is less than 3x for ONT data and 5x for WGBS data. The methylation frequency for each bin was calculated by averaging frequencies for positions assigned to the given bin.

Moreover, we compared the coverages and the number of calls for Rockfish and WGBS. We calculated strand-specific sequencing coverage for both ONT and WGBS by counting the total number of sequenced bases and dividing the number with two times the size of human genome (3.117 Gbp) cov = n_*bases*_/(2 × 3.117 × 10^9^). Complementary cumulative probability distribution for a specific coverage is defined as the proportion of CpG sites with equal or higher strandspecific calling coverage divided by the all CpG sites ccdf (cov) = *N*_≥*cov*_/*N*_*sites*_, ∀*cov* ∈ ℕ_0_. The calling coverage is the coverage provided by the final output for each tool. We define three types of highly confident CpG positive and negative sites: (1) sites with the calls concordant between both WGBS and Rockfish, (2) sites with the calls discordant between WGBS and Rockfish with the target method (WGBS or Rockfish) being supported either by Megalodon and/or Nanopolish and (3) sites for which either Rockfish or WGBS does not produce any call. A CpG site is defined to be positive if the strand-specific coverage is at least 5x with more than 50% methylation frequency. A CpG site is labelled as negative if the strand-specific coverage is at least 5x and methylation frequency is less or equal to 50%. A CpG site is deemed uncalled if coverage is less than 5x.

Lastly, we plotted the running time for every ONT tool. Experiments were repeated three times, with the bar sizes representing the average running time for each step. The experiments were run on the system with the following configuration: 2x AMD EPYC 7742 CPUs, 1TB DDR4 RAM, 8x NVIDIA A100 42GB VRAM GPUs with SSD storage. We limit each tool to 32 threads or processes and one GPU. Running time was computed using hyperfine^3^. Detailed commands are given in the Supplementary Material.

Rockfish was compared against two ONT-based methods, Megalodon (version 2.4.2) and Nanopolish (0.14.0). Guppy (5.0.14) supper accurate model was used as the canonical basecalling backend and Remora (0.1.2) was used as the modification calling backend in Megalodon. Internally Megalodon uses minimap2 (2.24; via Mappy) for pairwise alignment. For Nanopolish, reads were basecalled with Guppy (5.0.14) and aligned with minimap2 (2.24). For the WGBS methylation pipeline, we used Trim galore (0.6.7) to perform quality and adapter trimming. After trimming, reads were processed using Bismark (0.23.1). More details about the tools and examples of commands used for evaluation can be found in Supplementary Material.

### 4.4 Datasets

#### 4.4.1 Mouse datasets

Three mouse samples were used for the knowledge distillation training. C57BL/6 mice were bred and maintained under standard 12:12 h light/dark conditions at the National University of Singapore, by Miss Wang Xiao Jenny and Mr Li Yiqing Peter. The mice for cardiomyocyte isolation experiments were maintained at standard conditions. The mice for diet control were maintained in the following manner: co-housed male mice from mixed litters (n = 5 per cage) were provided with purified high fat/high sugar (sucrose) diet (45% fat DIO diet, TD.08811, Envigo) ad libitum after weaning (starting from 3-4 weeks of age after born). Eight weeks into the study, the diets of the mice were switched to a standard chow diet (2018, 18% Protein Rodent Diet, Envigo) for another eight weeks. Water was also provided ad libitum.

All animal protocols and experiments were approved by the Institutional Animal Care and Use Committee at the National University of Singapore.

Blood from the facial submandibular vein was collected in EDTA-coated micro-containers (Cat #:365974, BD). Following immediate centrifugation, at 4 °C for 15 min at 2000 rpm, blood cell pellets were collected and stored at -80 °C until analysis. Upon thawing, the red blood cell was burst by RBC Lysis Buffer: 100 mM Tris, pH 7.5, 0.2 mM EDTA buffer [65, 66]. Then the genomic DNA (gDNA) in the buffy coat was extracted as described below.

The neonatal and adult mouse cardiomyocyte isolation was carried out following the previous literature [67][68].

#### 4.4.2 H1ESc

H1 human embryonic stem cells (H1ESc) were grown by Dr Matias Autio Ilmari. H1ESc were maintained in mTeSR (Stemcell Technologies, 85850) on growth factor-reduced Geltrex (1:200 dilution, Thermo fisher, A1413202) coated plates at 37 °C with 5% CO_2_. Cells were dissociated using ReLeSR (Stemcell Technologies, 05872) for gDNA extraction.

Cell pellets were resuspended in 10 mM Tris pH 7.5 buffer and digested with 400 µg RNase A (Thermo Fisher, USA) at 37 °C for 30 mins. Then 0.5% SDS (final concentration) and 600 µg Proteinase K (Thermo Fisher) were added and incubated at 50 °C for 3 hours for digestion. The reaction mixture was purified by phenol-chloroform extraction following standard protocol, with UltraPureTM Phenol:Chloroform:Isoamyl Alcohol (Thermo Fisher) followed by one-time chloroform extraction to remove residual phenol. Then the native gDNA was precipitated by 2.5 volume of absolute ethanol and 10% volume of 3 M sodium acetate (pH 5.2, Ambion) following standard protocol, and washed once with 70% ethanol. The pellet was dried for 5 minutes and slowly hydrated at 4 °C, in 10 mM Tris pH 7.5 buffer for more than 72 hours.

The hydrated native gDNA extracted was quantified and quality checked by two independent researchers, and the library prep and sequencing were carried out by the Integrated Genomics Platform of Genome Institute of Singapore.

#### 4.4.3 Nanopore sequencing

Nanopore whole genome sequencing (WGS) libraries were constructed using the SQK-LSK109 kit (Oxford Nanopore Technologies) according to the manufacturer’s instructions.

DNA samples were optionally sheared to 15 kbp target size using the long hydropore on the Megaruptor (Diagenode) based on DNA size distribution assessed by pulsed-field gel electrophoresis on the Pippin Pulse (Sage Science).

Final ligation sequencing libraries were loaded on FLO-MIN106D, R9.4.1 flowcells, and sequenced on the GridION (Oxford Nanopore Technologies). Optional nuclease-flushes with flow-cell wash kit, EXP-WSH003 (Oxford Nanopore Technologies), and reloading of libraries were performed as required for optimal data generation.

#### 4.4.4 WGBS

The DNA used in whole genome bisulfite sequencing (WGBS) was first sheared by Covaris S2 (Covaris, USA) at 10% duty cycle, 2 × 40 s, intensity 5, cycle per burst 200 at 100 µL. Resulting DNA fragments were library prepped with NEBNext Ultra II DNA Library Prep Kit for Illumina (NEB, USA, E7645L) strictly following manufacturer’s instructions, using a methylated adaptor (E7536A). Without amplification post-library prep, the resulting library was purified and size-selected by 0.75X Ampure beads. Then the eluted DNA was bisulfite-converted with EZ DNA Methylation-Lightning Kit (Zymo, USA, D5030T) and TrueMethyl oxBS-Seq Module without oxidation step (Tecan, Switzerland) following the manufacturer’s instruction. The DNA obtained was amplified by PCR for 7-9 rounds with Q5U Hot Start High-Fidelity DNA Polymerase (NEB, M0515S), and purified by 0.8X Ampure beads. The samples were submitted to Macrogen (South Korea) for HiSeqX 150 bp paired-end sequencing according to standard Illumina cluster generation and sequencing protocols.

#### 4.4.5 Training datasets

For training, we used a high-quality ONT GM24385 dataset^4^ that contains both ONT reads and Illumina reads produced using reduced representation bisulfite sequencing (RRBS). GM24385 is a human B-lymphocyte cell line obtained from a white male. To produce a high-quality training dataset we first aligned basecalled ONT reads on the human reference using minimap2. We kept only the reads with exactly one alignment with a mapping quality of 60 or more. Reads aligned to chromosomes 2-21, X and Y were used for training. Reads aligned to chromosome 22 were used for validation and reads aligned to chromosome 1 were left for evaluation. Bismark coverage report used for extracting ground truth was generated by running the script provided in the dataset repository (see footnote 4). For base model training and validation we chose positions with RRBS coverage of 50x or more. Moreover, we removed all positions that are partially methylated. ONT examples from chromosomes 2-21, X and Y that have been discarded due to the aforementioned filters have been stored for knowledge distillation training. At last, we randomly sampled 100 million training examples to be used for training.

#### 4.4.6 Knowledge distillation datasets

After base model training, we performed knowledge distillation [69]. Knowledge distillation is a model compression method that results in similar or better performance compared to a teacher model while reducing the time needed for training and inference. The teacher model used for knowledge distillation is the trained Rockfish model trained on high-confident GM24385 data. To improve generalization and to introduce more biological diversity, we have sequenced three new mouse datasets used during distillation training.

To build a dataset for knowledge distillation, we sampled 300 million examples: 150 million examples from the previously filtered data, and 50 million examples from each of the three mouse datasets. Next, we performed inference on the sampled data. Probabilities obtained from the teacher model were used as probabilities for knowledge distillation. Moreover, we added 30 million examples from the base model training. In total, we had 330 million examples of knowledge distillation training. All samplings used to build the knowledge distillation dataset were stratified samplings - there were exactly 150 million modified and 150 million unmodified examples.

#### 4.4.7 Evaluation datasets

To show the robustness of our method, we performed an extensive evaluation of datasets corresponding to different cell lines. Alongside the aforementioned GM24385 chromosome 1, we also used three B-lymphocyte cell lines: NA12878 [70], NA19240 [71], HX1 [72], cancer cell line K562 [37] and the newly sequenced human embryonic stem cell H1ESc. Except for GM24385, we used all somatic chromosomes and chromosome X. For male samples (HX1, H1ESc) we also included chromosome Y. All reads were aligned to the CHM13.

## 5 Data Availability

Both ONT and Illumina RRBS paired-end data for GM24385 is available via AWS at https://labs.epi2me.io/gm24385-5mc/. ONT NA12878 dataset is available via AWS at https://github.com/nanopore-wgs-consortium/NA12878. ONT data for NA12940 is available upon request from Chaisson et al. [71]. ONT data for HX1 is available at NCBI under project PRJNA533926. ONT data for K562 is available at Gene Expression Omnibus (GEO) under the BioProject GSE173688.

WGBS paired-end data for NA12878 are available at ENCODE portal [73] (https://www.encodeproject.org/) under accession numbers ENCFF798RSS and ENCFF113KRQ (replicate 1), ENCFF585BXF and ENCFF851HAT (replicate 2). RRBS single-end data for NA12940 is available at ENCODE under accession numbers ENCFF000LZS (replicate 1) and ENCFF000LZT (replicate 2). WGBS paired-end data for HX1 is available at NCBI under the BioProject PRJNA301527. WGBS paired-end data for K562 is available at ENCODE under accession numbers ENCFF413KHN and ENCFF567DAI (replicate 1), ENCFF336KJH and ENCFF585HYM (replicate 2).

All newly sequenced data (ONT and WGBS for H1ESc, ONT for mouse datasets) are available at NCBI under BioProject PRJNA876781 (available upon publication).

ChIP-seq data for NA12878 are available at ENCODE under accession numbers ENCFF000ARV (replicate 1), ENCFF000ARP (replicate 2), ENCFF000ARK (control 1) and ENCFF000ARO (control 2).

Gene annotations (http://courtyard.gi.ucsc.edu/∼mhauknes/T2T/t2t_Y/annotation_set_v2/CHM13.v2.0.cat_liftoff_v2.gff3) and RepeatMasker annotations (https://t2t.gi.ucsc.edu/chm13/hub/t2t-chm13-v2.0/rmsk/rmsk.bigBed) are downloaded from UCSC Genome Institute (UCSC GI). GC content data (https://hgdownload.soe.ucsc.edu/hubs/GCA/009/914/755/GCA_009914755.4/bbi/GCA_009914755.4_T2T-CHM13v2.0.gc5Base.bw) and CpG island annotations are downloaded from UCSC Baskin School of Engineering.

## 6 Code Availability

All code, including feature extraction, model layout, training and inference can be found at https://github.com/lbcb-sci/rockfish. Both the base ^5^ and the small ^6^ model can be found on Google Drive. Moreover, they can be automatically downloaded using the script provided in the repository.

## 7 Acknowledgements

This work has been supported in part by Croatian Science Foundation under the project Single genome and metagenome assembly (IP-2018-01-5886), by Epigenomics and Epitranscriptomics Research seed grant from Genome Institute of Singapore (GIS), by Career Development Fund (C210812037) from A*STAR, and by the A*STAR Computational Resource Centre through the use of its high-performance computing facilities.

We would like to thank Dr Matias Autio Ilmari, Dr Matthew Andrew Ackers-Johnson, Ms Wang Xiao Jenny and Mr Li Yiqing Peter for their help with the cell and tissue samples used in this paper, and their helpful insight for this project. Moreover, thanks to Dr Yue Wan and Ms Sara Bakic for their valuable comments on the manuscript and data presentation. Finally, we would also like to thank Mr Low Hwee Meng and Ms Leong See Ting from GIS Integrated Genomics Platform core facility for their help and diligent work in Nanopore Sequencing.

## 8 Author Contributions

M.S. and L.Z. conceived the project. D.S. designed and implemented Rockfish with help from M.S and L.Z.; L.Z. performed ONT and WGBS sequencing of human data and ONT sequencing of mouse data. D.S. performed bioinformatics analysis and evaluation with the contribution of M.S. and L.Z.; D.S. and M.S. organized the manuscript. D.S., M.S. and L.Z. wrote the manuscript with help from R.F.; M.S supervised the project. M.S. and R.F. provided mentorship and support during the project.

## Footnotes

1 https://nanoporetech.com/nanopore-sequencing-data-analysis

2 https://github.com/FelixKrueger/TrimGalore

3 https://github.com/sharkdp/hyperfine

4 https://labs.epi2me.io/gm24385-5mc/

5 https://drive.google.com/file/d/1CXWcnKrrv9jJ3XirzZDV3tzW4MZyYj73/view?usp=sharing

6 https://drive.google.com/file/d/1-MJrnzknj2TIKzKSrT_UNfSooERdGc6Q/view?usp=sharing

