## Supplementary information for "Rockfish: A Transformer-based Model for Accurate 5-Methylcytosine Prediction from Nanopore Sequencing"

### Supplementary material

#### Running Guppy

Guppy is a basecalled developed by ONT. It is used for canonical base calling by methylation calling tools. The guppy version used in all experiments is 5.0.14, with a super accurate model. An example of the command is:

```
guppy_basecaller -i <fast5_folder> -r -s <output_folder> \
    --config dna_r9.4.1_450bps_sup.cfg --device <devices> \
    --fast5_out # Used for storing move table
```

#### Running Megalodon

Megalodon is a methylation-calling tool developed by ONT. It relies on canonical and modification basecalling provided by Guppy (in the recent versions Guppy is used only for canonical basecalling, while Remora is used for modification calling). It improves Guppy's (or Remora's) performance by anchoring basecalling output to a reference. First, a basecalled sequence is aligned onto the reference genome. Next, every candidate (5mC in CpG on a read-level) modification is evaluated against the canonical base using Viterbi scoring. The position is labelled as modified if the score corresponding to the modified base is higher than the score corresponding to the canonical base. Otherwise, the candidate is labelled as unmodified.
Megalodon version used for all experiments is 2.4.2, with Guppy (5.0.14) and Remora (0.1.2) used as backends. Minimap (v2.24) is provided via mappy. An example of a Megalodon command is:

```
megalodon \
    <fast5_folder> \
    --guppy-config dna_r9.4.1_450bps_sup.cfg \
    --guppy-server-path <guppy_basecall_server_path> \
    --remora-modified-bases dna_r9.4.1_e8 sup 0.0.0 5mc CG 0 \
    --outputs per_read_mods mods \
    --write-mods-text \
    --reference <reference_fasta> \
    --devices <gpus> \ # format: cuda:d_0,d_1,… (e.g. cuda:0,1,2)
    --processes <n_processes>
```

#### Running Nanopolish

Nanopolish is a methylation calling tool based on the hidden Markov model (HMM). First, basecalled reads (fastq) are aligned using minimap2 (v2.24). Next, the raw nanopore signal corresponding to the aligned read is aligned to the reference sequence (event_align). For every candidate group (a region around one or multiple CpGs) Nanopolish calculates the ratio between the likelihood of methylation and the likelihood of unmethylation. If the ratio > 0, the region is labelled as methylated, otherwise as unmethylated.
Nanopolish version used for all experiments is 0.14.0. Commands used for running Nanopolish are given here:

1. Nanopolish index:
   ```
   nanopolish index -d <fast5_folder> <reads>
   ```

2. Nanopolish align:
   ```
   minimap2 -t <threads> -a -x map-ont <reference> <reads> | samtools
   sort -T tmp -o <bam>
   ```

```
samtools view -@ <threads> -F 2308 -bS <in_bam> > <out_bam>
samtools index -@ <threads> <bam>
```

3. Nanopolish call methylation:

```
nanopolish call-methylation -r <reads> -b <bam> \
    -g <reference> -q cpg -s reference \
    -t <threads> --min-mapping-quality 0 > <tsv_output>
```

Site-level frequency was calculated using the script available at:
https://github.com/jts/nanopolish/blob/v0.14.0/scripts/calculate_methylation_frequency.py
Since Nanopolish calls methylation for a region (not for individual position), all read-level calls are extracted using the algorithm that can be found in the code above (lines 52-67).

#### Rockfish installation and inference

Rockfish can be installed by cloning the GitHub repository (#TODO Add repo) and invoking pip install command:

```
git clone ... rockfish && cd rockfish
pip install --extra-index-url https://download.pytorch.org/whl/cu113 .
```

User should use replace “cu113” (in the link) with the desired CUDA version (e.g. for cuda 10.2 link is https://download.pytorch.org/whl/cu102).

Rockfish model can be downloaded using the command:

```
rockfish download -m <{all, base, small}> -s <save_path>
```

The argument “`all`” will download both the base and small models.

An example of Rockfish inference:

```
rockfish inference -i <basecalled/workspace> --reference <reference> \
    --model_path <model_path> -r -t <n_workers> \
    -b <batch_size> \  # Total batch size (all GPUs), default 8092
    -d <devices>  # Format: d_0,d_1,... (e.g.) "0,1,2"
```

Note: The input to the inference is workspace folder that can be found in the folder saved by Guppy. Workspace folder contains fast5 files annotated with fastq and move table.

#### WGBS pipeline

The pipeline for processing bisulfite data is similar to the pipeline given by ONT (https://ont-open-data.s3.amazonaws.com/gm24385_mod_2021.09/bisulphite/fastq2bed.sh). The pipeline consists of five steps:

1. Adapter and quality trimming using trim_galore
2. Alignment using bismark
3. Deduplication using deduplicate_bismark
4. Methylation extraction using bismark_methylation_extractor
5. Converting output from 4) to bedGraph using bismark2bedGraph

In the first step we use “`--three_prime_clip_R1 15 --clip_R2 15`” arguments only for GM24385 dataset. Moreover, deduplication (third step) was skipped for NA19240 since it is recommended to skip deduplication for RRBS (https://github.com/FelixKrueger/Bismark/issues/234).

### Supplementary Figures

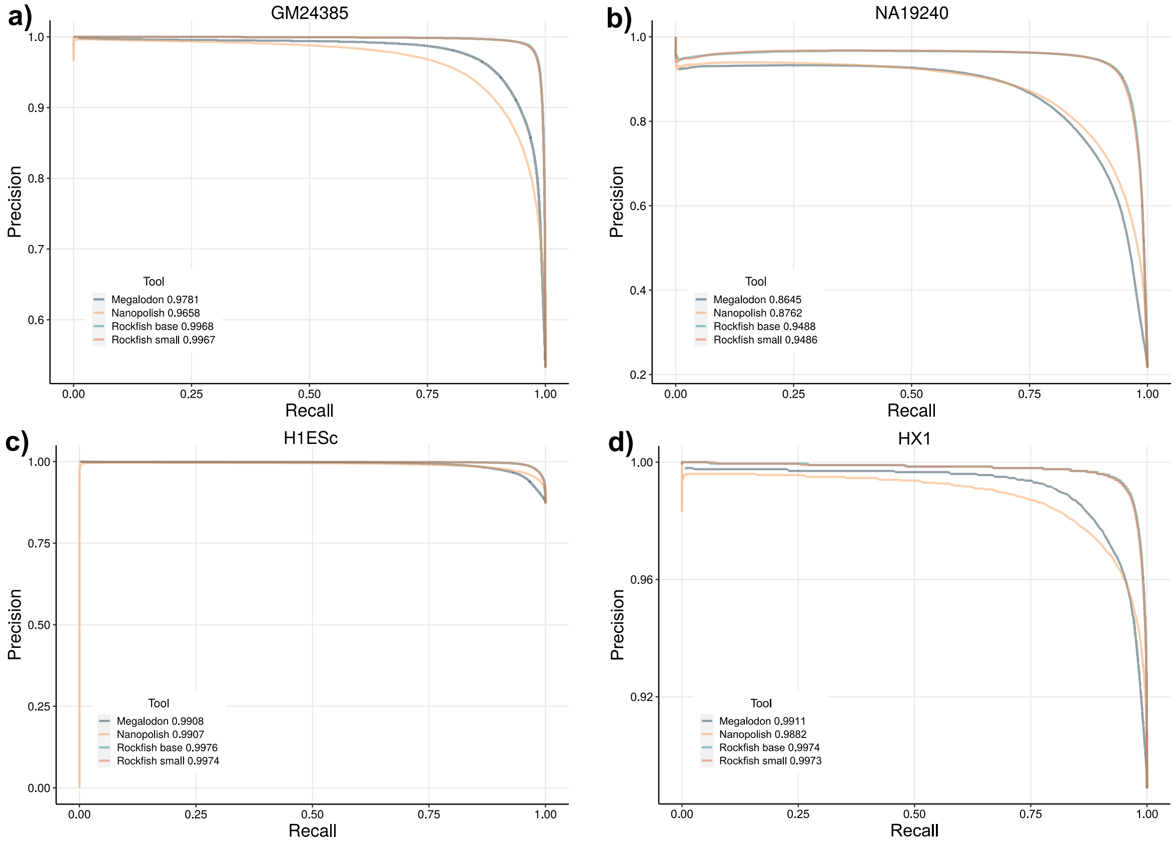


*Figure S1: Precision-recall curves on the read-level evaluation for **a)** GM24385, **b)** NA19240, **c)** H1ESc and **d)** HX1 datasets. Only examples predicted by every ONT tool are included. Included predictions are intersected with WGBS data. Partially methylated positions and positions with coverage less than x5 (WGBS) are excluded. Rockfish and Megalodon examples are sorted using modification probabilities. Nanopolish is sorted using the log-likelihood ratio value.*

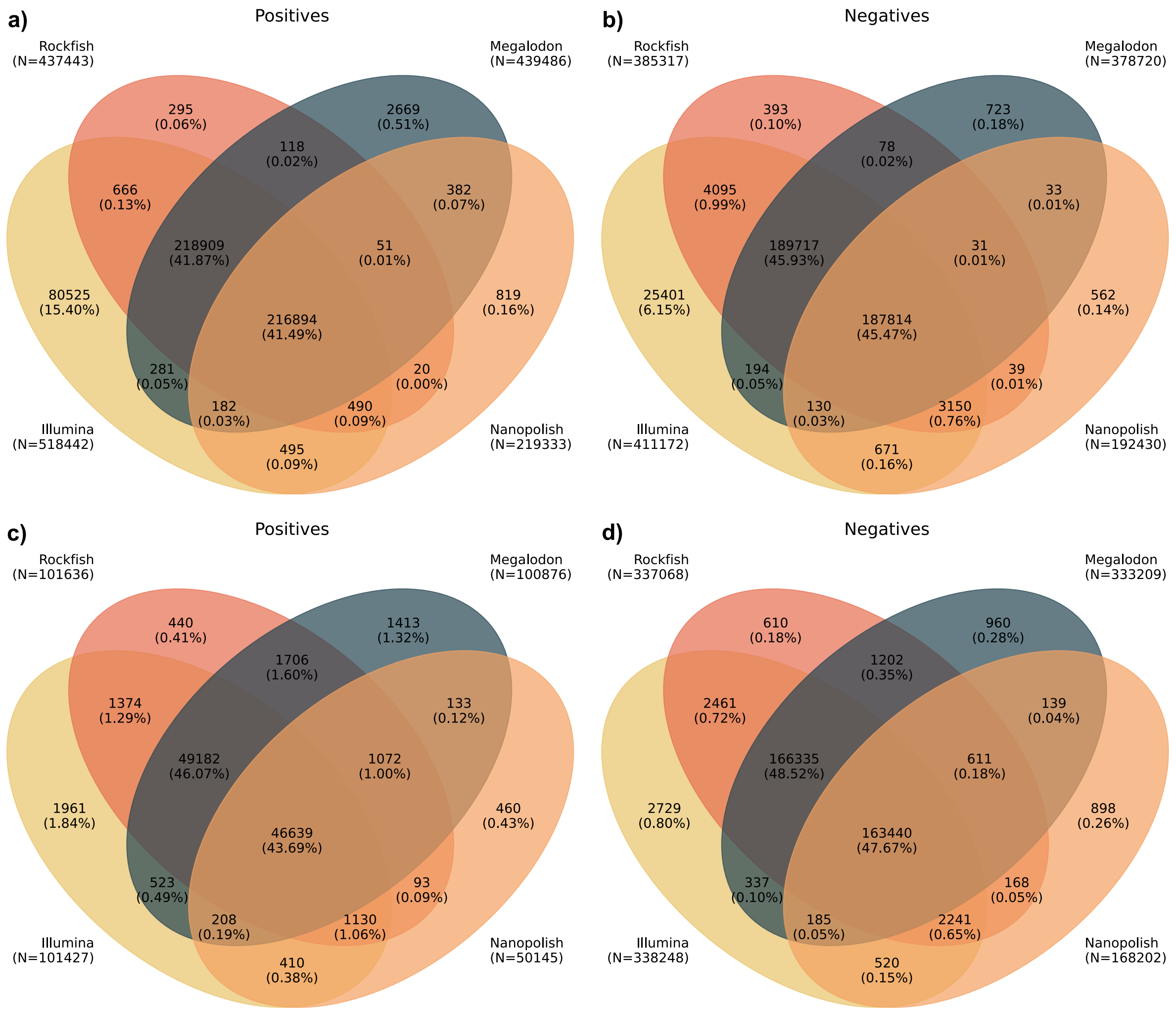


*Figure S2: Venn diagrams representing predicted positives and negatives for GM24385 (**a-b**) and NA19240 (**c-d**) datasets. Actual positives and negatives (the ground truth) are given by the set named "Illumina". Rockfish is represented with the small model. Sample space is defined as the set of all fully unmethylated or methylated sites called by Illumina with at least 5x. Rockfish calls the highest amount of true positives and true negatives and achieves high precision and recall.*

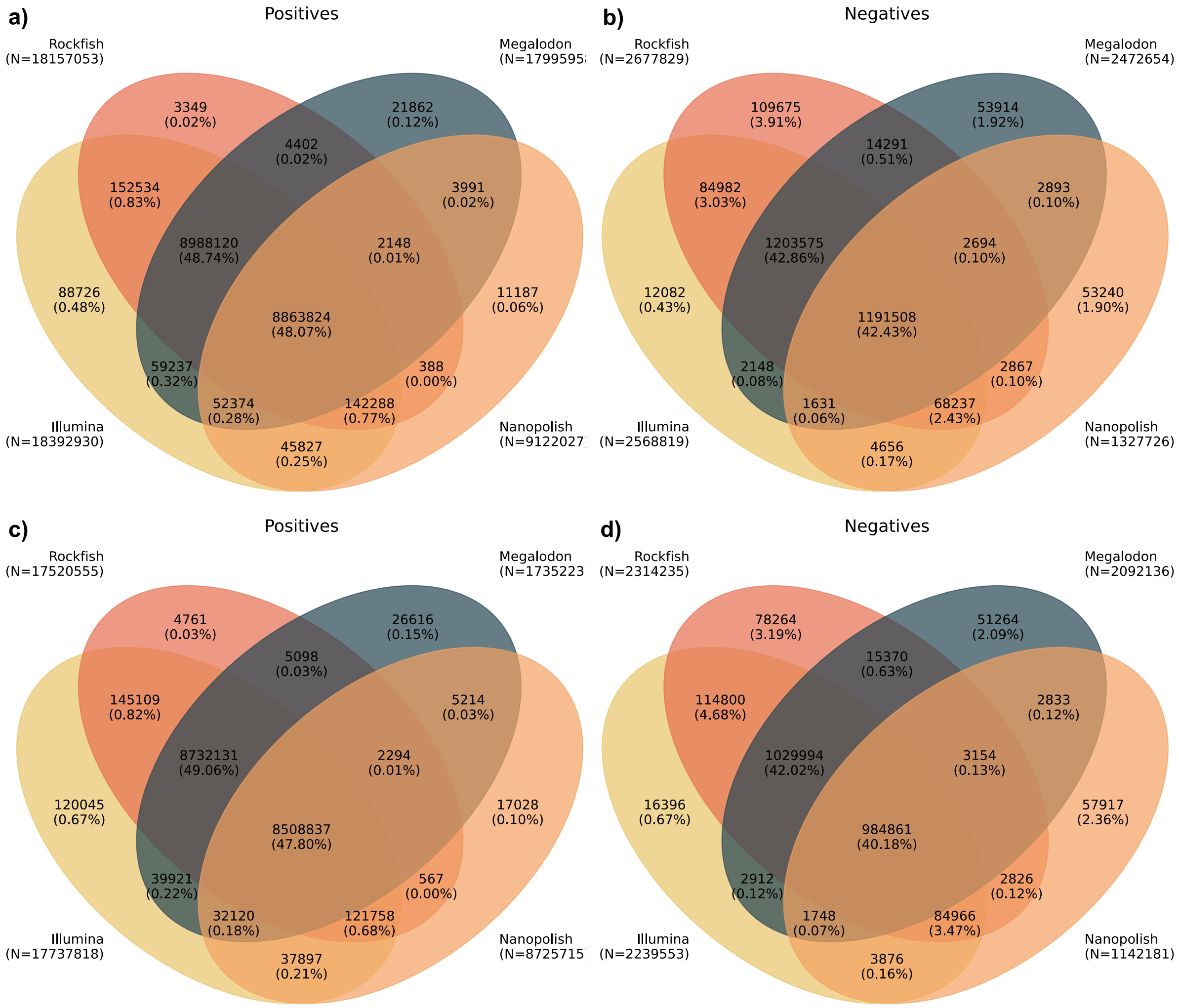


*Figure S3: Venn diagrams representing predicted positives and negatives for H1ESc (**a-b**) and HX1 (**c-d**) datasets. Actual positives and negatives (the ground truth) are given by the set named "Illumina". Rockfish is represented with the small model. Sample space is defined as the set of all fully unmethylated or methylated sites called by Illumina with at least 5x. Rockfish calls slightly more false negatives compared to Megalodon and Nanopolish, but significantly more true negatives.*

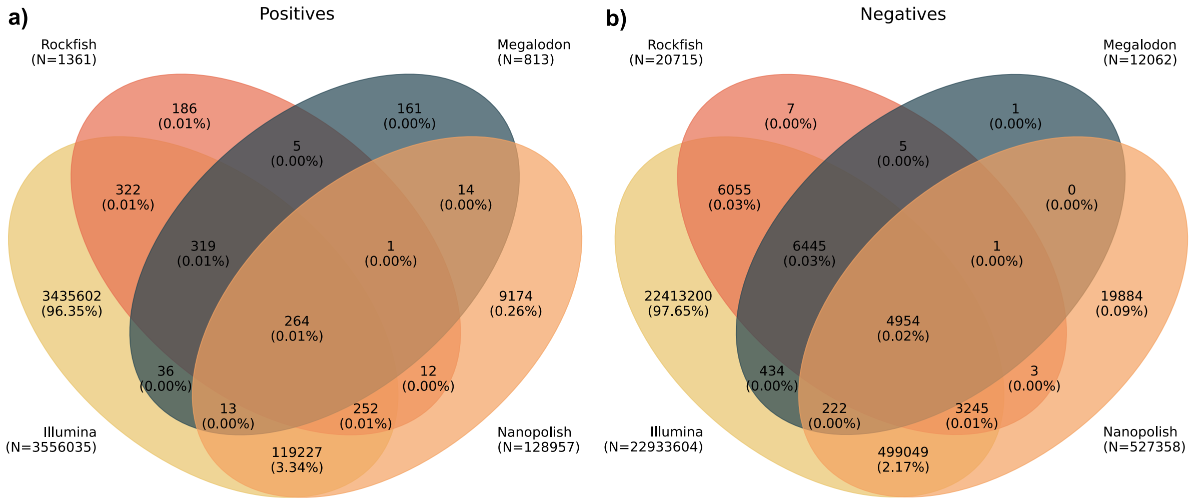


*Figure S4: Venn diagrams representing predicted positives (**a**) and negatives (**b**) for the K562 dataset. Actual positives and negatives (the ground truth) are given by the set named "Illumina". Rockfish is represented with the small model. Sample space is defined as the set of all fully unmethylated or methylated sites called by Illumina with at least 5x. An overwhelming majority of examples are not called by the ONT methods due to low ONT coverage.*

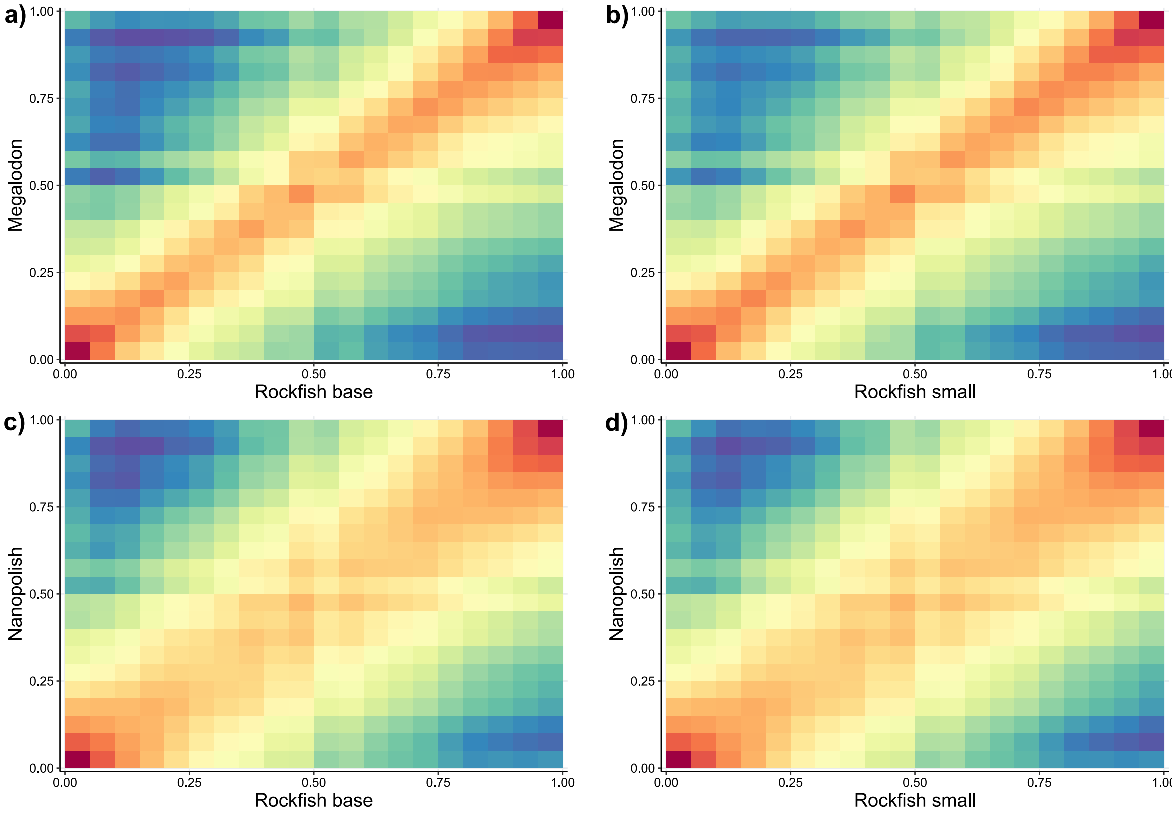


*Figure S5: 2D histograms showing the correlation between Rockfish models and other ONT tools for the NA12878 dataset. Each axis is divided into 20 bins, with counts being plotted on a log scale. Subfigure a) shows correlation between Rockfish base model and Megalodon (Pearson's r = 0.9495), b) Rockfish small and Megalodon (r = 0.9529), c) Rockfish base and Nanopolish (r = 0.9215) and d) Rockfish small and Nanopolish (r = 0.9240). P-value = 0 for all tests.*
